# Nuclease proteins CifA and CifB promote spermatid DNA damage associated with symbiont-induced cytoplasmic incompatibility

**DOI:** 10.1101/2022.04.04.487029

**Authors:** Rupinder Kaur, J. Dylan Shropshire, Brittany A. Leigh, Seth R. Bordenstein

## Abstract

The worldwide endosymbiosis between arthropods and *Wolbachia* bacteria is an archetype for reproductive parasitism. This parasitic strategy rapidly increases the proportion of symbiont-transmitting mothers, and the most common form, cytoplasmic incompatibility (CI), impacts insect evolution and arboviral control strategies. During CI, sperms from symbiotic males kill embryos of aposymbiotic females via two nuclear-targeting proteins, CifA and CifB, that alter sperm chromatin organization in *Drosophila melanogaster*. Here we hypothesize that Cif proteins metabolize nucleic acids of developing sperm to initiate genome integrity changes. Using *in vitro* and *in situ* transgenic, mutant, enzymatic, and cytochemical assays, we show that CifA is a previously-unrecognized DNase and RNase, and CifB is a DNase. Notably, *in vitro* nuclease activity translates to *in situ* spermatid DNA damage at the canoe stage of spermiogenesis. Evolution-guided mutations ablate Cif enzymatic activity. Nucleic acid metabolism by Cif enzymes expands a fundamental understanding of the mechanism of symbiont-mediated reproductive parasitism.

## Introduction

*Wolbachia* are maternally inherited, obligate intracellular bacteria that occur worldwide in 40-65% of arthropod species (Hilgenboecker et al., 2008; Kaur et al., 2021; Weinert et al., 2015; Zug and Hammerstein, 2012). Many *Wolbachia* strains selfishly alter host reproduction to increase the relative number of symbiotic females that transmit the bacteria to the next generation (Hoffmann et al., 1990; Hoffmann and Turelli, 1997; Kriesner et al., 2013). Cytoplasmic incompatibility (CI) is the most studied reproductive phenotype with major impacts on arthropod evolution and vector control. Specifically, CI results in embryonic death when symbiotic males mate with aposymbiotic females. Nullification of death, and thus rescue of CI, occurs when transmitting females harbor the same strain of *Wolbachia* (Shropshire et al., 2020). CI-causing mosquitoes are released in both population suppression (Caputo et al., 2020; Mains et al., 2016; Puggioli et al., 2016; Zheng et al., 2019) and population replacement strategies (Flores and O’Neill, 2018; Indriani et al., 2020; Nazni et al., 2019; Ryan et al., 2020) to curb arbovirus transmission of Dengue, Zika, and Chikungunya (Caragata et al., 2016; Crawford et al., 2020; Dobson et al., 2002; Hoffmann et al., 2011; Moreira et al., 2009; O’Neill, 2018; O’Neill et al., 2018). Thus, CI is at the forefront of controlling mosquito-borne diseases globally (Bourtzis et al., 2014; O’Neill et al., 2018; Utarini et al., 2021).

CI is caused by the germline expression of two cytoplasmic incompatibility factor genes, *cifA* and *cifB,* in *w*Mel-infected *Drosophila melanogaster* testes (Beckmann et al., 2017; LePage et al., 2017), and *cifA* expression in ovaries rescues CI (Shropshire et al., 2018). Thus, CI is governed by the Two-by-One genetic model demonstrated in various systems (Beckmann et al., 2019; Chen et al., 2019; LePage et al., 2017; Shropshire et al., 2021b; Shropshire and Bordenstein, 2019). *cifB* alone can also cause *cifA*-rescuable CI under a One-by-One genetic model (Adams et al., 2021; Sun et al., 2021). Therefore, deciphering the Cif molecular mechanism(s) and potentially universal features of CI and rescue remains a central goal for understanding how sexual reproduction serves as a battleground for host-*Wolbachia* interactions.

CifA and CifB proteins from *w*Mel of *D. melanogaster* and *w*Pip of *Culex pipiens* were recently found to invade the nuclei of developing sperm (Horard et al., 2022; Kaur et al., 2022). Specifically, dual expression of *w*Mel CifA and CifB alters sperm chromatin integrity before fertilization via modulating the histone-protamine transition of spermiogenesis (Kaur et al., 2022), and *w*Pip CifB expression impacts sperm DNA stress in the embryo (Horard et al., 2022). Sperm nuclear access by *w*Mel Cifs indicates that Cifs may directly interact with sperm DNA and/or RNA to modify sperm genome integrity and contribute to the CI mechanism. Thus, we hypothesized that Cifs may metabolize sperm nucleic acids by their enzymatic properties.

Enzymatic functions of CifA have not been tested, while *in vitro* enzymatic properties of some of the CifB variants have been evaluated. T1 *w*Pip CifB is a deubiquitinase (DUB) that cleaves poly-ubiquitin chains *in vitro* to cause CI (Beckmann et al., 2017). However, mutating the catalytic residue in CifB’s DUB domain ablated CI in one study (Beckmann et al., 2017) and maintains CI in another (Horard et al., 2022), suggesting that DUB enzymatic function may not be essential for CI. T4 *w*Pip CifB is a DNase with active PDDEXK nuclease domains that cleave DNA substrates (Chen et al., 2019). Presumptive activation of PDDEXK nuclease sites with artificial amino acid substitutions in T1 *w*Pip CifB did not degrade DNA *in vitro* (Chen et al., 2019), suggesting these domains are inactive in T1 CifB variants. Notably, the enzymatic activity of wild-type T1 *w*Mel CifB remains untested to date. Mutating conserved amino acid residues in the nuclease and other domains of *w*Mel CifB ablates CI (Shropshire et al., 2020), suggesting that nuclease domains and various other sites throughout CifB are crucial for CI, thus warranting *in vitro* and *in situ* enzymatic characterization.

CifA is weakly predicted to encode a domain of unknown function (DUF3243) with homology to a Puf-family RNA-binding domain found throughout eukaryotes, and mutating conserved residues in this domain notably ablates CI but not rescue (Shropshire et al., 2020). In contrast, mutating the conserved residues of the sterile-like transcription factor (STE) domain of CifA had no impact on CI (Shropshire et al., 2020), establishing the hypothesis that these domains and residues may possess a molecular function relevant to CI. Moreover, a nuclear localization sequence is necessary for CifA-mediated CI, rescue and maintaining sperm genome integrity (Kaur et al., 2022), thus motivating interest to understand CifA’s enzymatic roles in nucleotide metabolism.

Here, we performed *in vitro* and *in situ* biochemical assays with wild type, transgenic, and mutant proteins to determine the nuclease capabilities of T1 *w*Mel CifA and CifB proteins. We demonstrate that (i) both CifA and CifB are *in vitro* nucleases, where CifA cleaves single-stranded (ss) DNA, double-stranded (ds) DNA and ssRNA substrates, and CifB cleaves both DNA substrates, (ii) Evolutionary-guided substitutions and truncations ablate nuclease functions, (iii) *w*Mel *Wolbachia* and transgenic *cif* expression causes *in situ* spermatid DNA fragmentation in *D. melanogaster*. We discuss the importance of these findings to the mechanism of the Cif proteins in inducing CI.

## Results

### Type 1 CifB protein is an *in vitro* DNase

To test nuclease activity of T1 *w*Mel CifB, we first generated CifB recombinant proteins for *in vitro* enzyme assays against DNA and RNA oligonucleotide substrates. The full CifB protein is too large to recombinantly express in *E. coli* (Wang et al., 2022; Xiao et al., 2021). As such, we generated proteins spanning the unannotated amino terminus (A), the N-terminal nuclease domain (NTND) and C-terminal nuclease domain (CTND) (labeled as CifB_△D_, 1-796 aa), both nuclease domains only (CifB_△A△D_, 277-796 aa) (**Fig. 1A**), and A-terminus alone (CifB_△;N△C△D_, 1-276 aa) (**Fig. S1**). We previously demonstrated that mutations in highly conserved residues across CifB, including both nuclease domains (labeled CifB_2_ and CifB_3_), ablate transgenic CI (Shropshire et al., 2020), suggesting these domains are crucial for the CI mechanism. Therefore, we generated CifB recombinants with the same substitutions in the NTND (CifB_2;△D_) and CTND (CifB_3;△D_) (**Fig. 1A**) to test if these same mutants ablate nuclease activity *in vitro*.

**Figure 1.**
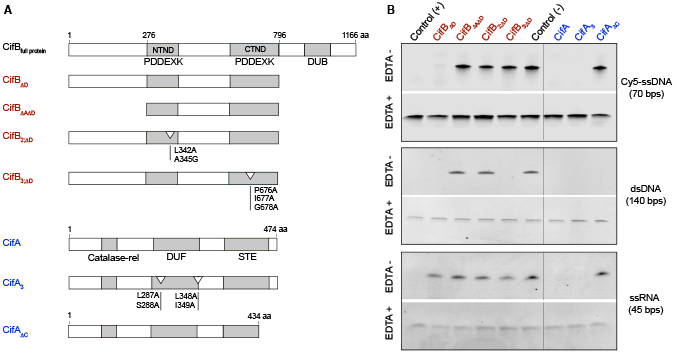
CifA and CifB proteins are *in vitro* nucleases in which CifA is a DNase and RNase, and CifB is a DNase. (A) Schematic representation showing the Cif recombinant proteins used in the study. Since the full CifB protein is too large to be recombinantly expressed in *E. coli,* purified His-GST tagged CifB variants were generated with a mixture of the amino terminus, N-terminal nuclease domain (NTND), and C-terminal nuclease domain (CTND) along with various engineered substitutions in conserved residues. The full His-tagged CifA protein was generated with and without substitutions in conserved residues and a truncation in the C-terminus. (B) Nuclease activities of the Cif proteins against DNA and RNA substrates. CifB shows both ss- and ds-DNase activity. While ss-DNase activity is ablated by mutations in both NTND (CifB_2;△D_) and CTND (CifB_3;△D_), ds-DNase activity is ablated by substitutions in NTND but not in CTND. CifA is both a DNase and RNase. Substitutions in conserved residues of the DUF domain (CifA_3_) do not alter CifA nuclease activity, while a truncation at the C-terminus (CifA_△C_) ablates ss-DNase and RNase action of CifA. DNase and RNase enzymes were used as positive controls, and no CifA or CifB proteins were added to the reaction mixtures for negative controls. EDTA was added to a 20x molar excess over Mg^2+^ to inhibit the reactions. Cy5-labeled ssDNA samples were run in a 10% polyacrylamide/TBE gel. Non-labeled dsDNA and ssRNA samples were run in 10% polyacrylamide/TBE gel and stained with GelRed. Assays were replicated three times independently. Full uncropped gel images are submitted in source data files. Related reagents can be available upon request from Bordenstein lab.

We report four key outcomes:

First, CifB_△D_ with the amino terminus and both nuclease domains cleaved ssDNA and dsDNA substrates similar to the control DNase enzyme (**Fig. 1B**). As expected, ethylenediaminetetraacetic acid (EDTA) ablated the nuclease activity since it chelates divalent cations required for nuclease activity. We, therefore, conclude that T1 CifB_△D_ (referred to as CifB hereafter) encodes DNase enzymatic function, which is contrary to previous claims that tested recombinant protein with presumptive mutations thought to recapitulate catalytic activity (Beckmann et al., 2017; Chen et al., 2019).

Second, the truncated CifB without the amino terminus (CifB_△A△D_) failed to cleave ssDNA or dsDNA, suggesting the amino terminus plays a crucial role in nuclease activity. The amino terminus, for instance, may affect proper structural folding of the protein or binding to the target substrate. The amino terminus alone (CifB_△N△C△D_) also did not exhibit cleaving activity (**Fig. S1**), reinforcing the importance of the intact amino terminus together with both nuclease domains. Intact CifB’s DNase activity was substantiated once more against a ssDNA substrate using a range of protein concentrations and reaction incubation periods in which enzymatic activity decreases with lower enzyme concentration and shorter incubation time (**Fig. S2**), as expected.

Third, mutant CifB proteins with substitutions in each nuclease domain (CifB_2;△D_ and CifB_3△D_) that ablate transgenic CI in flies (Shropshire et al., 2020) also abolished ssDNA cleavage here *in vitro* (**Fig. 1B**). Findings suggest that substitutions in these negatively-selected residues (Shropshire et al., 2020) are important for the ss-DNase activity of CifB. Moreover, incubation of these proteins with dsDNA revealed that CifB_2△D_ ablates ds-DNase activity, whereas CifB_3△D_ did not.

Fourth, we investigated if CifB is a RNase and found that CifB variants did not cleave ssRNA oligonucleotide substrates (**Fig. 1B and SI**). We also performed mass spectrometry of the purified CifB protein to rule out the alternative explanation that co-purified *E. coli* nucleases with the CifB protein confound the CifB nuclease activity observed. We detected no contaminating DNase enzymes (**Table S1**).

Taken together, CifB is a DNase, and the CifB_2_ sites in the NTND are crucial for both ssDNA and dsDNA cleavage, whereas CifB_3_ sites in CTND are substrate-specific as the mutated residues only ablate cleavage of ssDNA. These data are semi-consistent with crystal structures of Cif proteins (Xiao et al., 2021) that showed the NTND of T4 CifB from *w*Pip strain is the main catalytic center, and the CTND may be involved in specific substrate binding.

### Type 1 CifA protein is an *in vitro* DNase and RNase

Purified full-length CifA was incubated with the same DNA and RNA substrates as above. We report that CifA cleaved the ssDNA, dsDNA, and ssRNA substrates in the absence of EDTA (**Fig. 1B, Fig. S2**). Notably, CifA’s nuclease activity is never reported before. Mass spectrometry of the CifA protein once again indicated there was no co-purified DNase from *E. coli* that could alternatively explain CifA’s DNase function (**Table S1**). However, there was a very low abundance of ribonuclease peptides from *E. coli* in the purified CifA construct (**Table S1**). To test if the *E. coli* RNase contaminant contributed to CifA’s RNase activity, we performed nuclease activity assays by incubating RNA substrate with various concentrations of CifA protein diluted up to 16-fold to eliminate the rare contaminant. CifA RNase activity persisted upto 8-fold enzyme dilutions (**Fig. S2A**). Moreover, the same low abundant *E. coli* RNase contaminant was also detected in the purified CifB sample (**Table S1**) that notably lacked RNase activity. These results support the RNase properties of CifA and that the minute traces of randomly distributed *E. coli* contaminants are irrelevant. CifA’s RNase function is consistent with predictive CifA annotations for RNA binding and transcription.

CifA has low sequence similarity to a domain of unknown function (DUF3243) with structural homology to a RNA-binding domain (Lindsey et al., 2018). We previously showed that mutating conserved residues in the T1 CifA DUF (labeled CifA_3_) ablated CI but not rescue in transgenic flies (Shropshire et al., 2020). Thus, we tested here whether CifA_3_ (**Fig. 1A**) cleaves DNA and/or RNA. CifA_3_ retained nuclease activity (**Fig. 1B**), suggesting that the conserved residues in the DUF are crucial to CI but do not contribute to CifA’s *in vitro* nuclease activity. In contrast, CifA_△C_ protein with a 40 amino acid truncation inclusive of the predicted STE transcription factor at the C-terminus ablated ss-DNase and ss-RNase function, but dsDNA cleavage remained (**Fig. 1B**). These findings suggest that the C-terminus of CifA contributed to substrate specificity for ssDNA and ssRNA. Notably, this region is proposed to be disordered based on the crystal structure of CifA (Wang et al., 2022; Xiao et al., 2021). Disordered regions can regulate ssDNA binding in *Escherichia coli* (Kozlov et al., 2015), facilitate post-translational modifications, and often regulate key protein functions (Sharma and Schiller, 2019).

In *D. melanogaster,* dual expression of *w*Mel CifA with CifB is required to induce CI (Beckmann et al., 2017; Kaur et al., 2022; LePage et al., 2017; Shropshire and Bordenstein, 2019). To examine if T1 *w*Mel CifA and CifB have any impact on their independent nuclease activities, we co-incubated the proteins with dsDNA or ssRNA substrates as described earlier (Chen et al., 2019). We found that DNA and RNA cleavage continued upon co-incubation of CifA variants with different active and inactive CifB variants **(Fig. 2)**.

**Figure 2.**
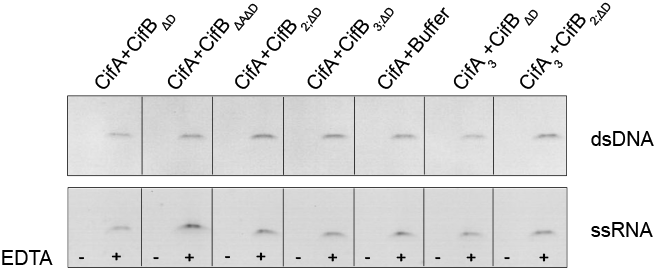
Nuclease activity persists when CifA and CifB variants are co-incubated. Prior to adding enzymes to the dsDNA and ssRNA substrates, 10 μM of CifA and 1 μM CifB variants were co-incubated on ice for 30 min to allow complex formation. To serve as a negative control, CifA was also co-incubated with the storage buffer only, in which CifB purified proteins were stored. The reactions were carried out for 120 min. EDTA was added to a 20× molar excess over Mg^2+^ to inhibit the reactions. Samples were run in 10% polyacrylamide/TBE gel and stained with GelRed. Full uncropped gel images are submitted in source data files. Related reagents can be available upon request from Bordenstein lab.

### wMel *Wolbachia* and transgenic Cif expression promotes *in situ* spermatid DNA damage

We next sought to evaluate if *in vitro* Cif nuclease activity translates to *in situ* activity by dissecting whole testes from <8 hrs old wild type (*w*Mel+ and *w*Mel-) and transgene-expressing *Drosophila* males. We performed TUNEL (terminal deoxynucleotidyl transferase dUTP nick-end labeling) assays to examine DNA breaks in spermatid nuclei isolated from testes squashes. DNA breaks and repair are typical features accompanying the histone-to-protamine transition during the canoe stage of spermiogenesis to tightly pack the chromatin in the sperm nucleus (Rathke et al., 2007). However, excessive DNA breaks cause sperm chromatin abnormalities and adversely impact male fertility status in mammals (Agarwal and Said, 2003). Therefore, we investigated CifA- and CifB-induced spermatid DNA damage with TUNEL signals upon transgenic and wild-type expression of the Cifs relative to negative controls.

TUNEL signals occured in the elongating spermatids at the canoe stage and were no longer detected in the individualized and mature sperms in all treatment groups, as expected (Rathke et al., 2007) (**Fig. S3**). In both CI-inducing *w*Mel and *cifAB*-expressing males, there were significantly higher percentages of sperm bundles with strong TUNEL intensity relative to their aposymbiotic controls (**Fig. 3**; Mann-Whitney test, p<0.0001, **Table S2**), indicating the Cifs caused the increased DNase activity. CifB initiated significantly higher DNA damage in single expressing lines than CifA and negative control lines (**Fig. 3**; Kruskal-Wallis test, p<0.0001, **Table S2**), suggesting CifB with and without the DUB domain can cleave DNA *in situ* and *in vitro,* respectively. Notably, transgenic expression of individual CifB from *w*Mel does not cause CI in *D. melanogaster* (**Fig. S4, Table S3**) (LePage et al., 2017; Shropshire and Bordenstein, 2019). CifB-mediated *in situ* DNase activity alone is therefore insufficient to induce CI. CifA did not cause *in situ* DNase activity, which contrasted with the *in vitro* data above. We hypothesize that CifA’s ability to cleave spermatid nuclear DNA may depend on the co-expression of CifB, or CifA’s RNase activity in the testes (not observable by TUNEL here) may be relevant to CI (**Fig. 5**).

**Figure 3.**
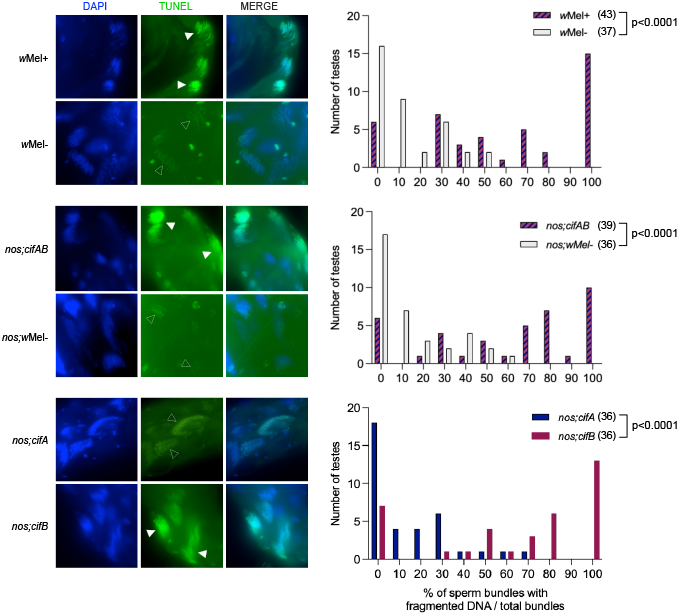
*w*Mel *Wolbachia* and transgenic Cif expression cause *in situ* DNase activity in canoe-stage spermatids. TUNEL staining on testes squashes from <8 hrs old males was performed to visualize and quantify sperm bundles with DNA breaks across the genotypes tested. Compared to the negative controls *w*Mel-(empty arrow heads), DNA breaks marked by TUNEL staining (green) and DNA (blue) are abundant in the wild type *w*Mel *Wolbachia* and dually expressed *cifAB* transgenic lines at the canoe stage of spermiogenesis (solid arrow heads). Upon individual expression, CifB induces DNA damage, while CifA does not. Total sperm bundles and those with fragmented DNA were manually counted. The numbers of testes investigated are shown in parentheses next to the genotype. The experiment was performed with two independent biological replicates and samples were blind-coded for the first run. P-value significance was calculated by Mann-Whitney pairwise comparison test. P-values are reported in Table S2. CI-based hatch rate assay was conducted in parallel as shown in Figure S4.

**Figure 4.**
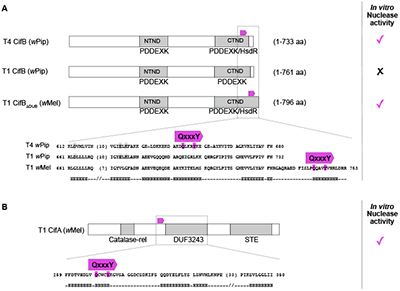
Nuclease activity of Type 4 CifB and Type 1 CifA and CifB associates with a QxxxY motif. (A) Schematic of CifB proteins and amino acid sequence showing the CTND location of QxxxY motifs that belong to the HsdR subunit of RecB nuclease family (Singleton et al., 2004). The motif and glutamine (Q) and tyrosine (Y) residues are highlighted in magenta, whereas the previously characterized aspartate (D), glutamate (E) and lysine (K) residues belonging to the PDDEXK nuclease family (Chen et al., 2019) are highlighted in grey. The QxxxY motif is present in *w*Pip Type 4 (T4) CifB known as an *in vitro* nuclease (Chen et al., 2019) and T1 CifB from *w*Mel *Wolbachia* characterized as an *in vitro* nuclease (this study). The QxxxY motif is absent in the non-nuclease *w*Pip T1 CifB. (B) T1 CifA protein from *w*Mel *Wolbachia* shown to be an *in vitro* nuclease (this study) also contains a QxxxY motif in a predicted α-helical region. Predicted α-helical residues are labeled “H” and residues predicted to be part of β-sheets are labeled “E.” The numbers of excluded residues are shown in parentheses. The last residue numbers are shown at the end of each sequence.

**Figure 5.**
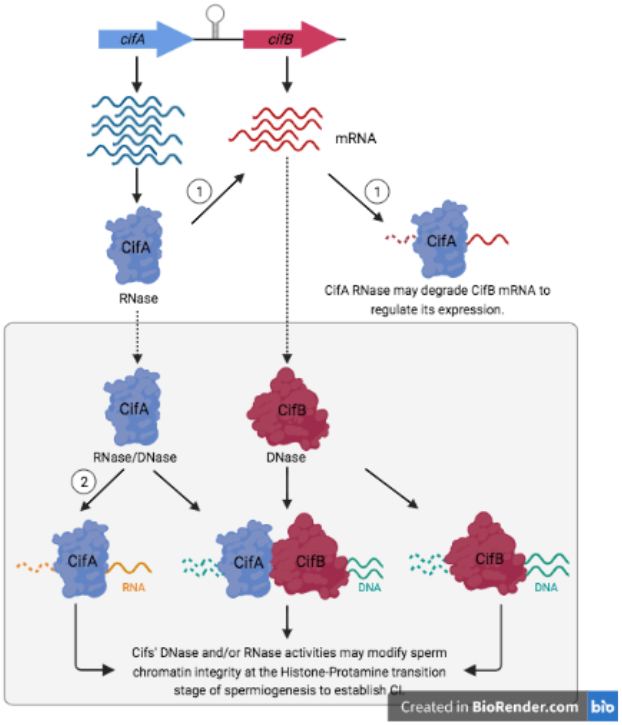
Illustrative summary and hypotheses on CifA and CifB nuclease functions under the Host Modification model of CI. *cifA* and *cifB* genes are shown in blue and red arrows, respectively, with an annotated hairpin terminator. *cifA* is highly transcribed compared to *cifB* (Kaur et al., 2022). CifA enzyme encodes *in vitro* DNase and RNase activities. 1. Under the type V toxin-antitoxin system (52), the CifA RNase may degrade mRNA of the *cifB* gene and suppress its toxicity against the host. 2. Under the Host Modification model of CI, CifA RNase may target spermatogenesis-associated mRNA(s) to modify sperm genome integrity and induce CI. CifB’s DNase activity alone or combined with CifA’s DNase and/or RNase functions may induce DNA fragmentation in canoe spermatids, the stage when histones are replaced by protamines to generate tightly condensed chromatin (Rathke et al., 2014). CifB’s DNase action is necessary for CI in some systems (30), though the DNA substrate in the host remains uncharacterized. On the other hand, we conversely show here that CifB spermatid DNase activity from *w*Mel *Wolbachia* is insufficient to cause CI, which may be contingent upon CifA’s RNase and/or DNase activity that could hypothetically modify sperm genome integrity. CifAB-induced sperm modifications at this stage may pave the incipient steps to establish CI.

Most, but not all, of the conserved residues that were mutated in CifA and CifB ablated or decreased CI strength in flies (Shropshire et al., 2020). To check if these same variants contribute to *in situ* DNA damage of the developing spermatids, we first performed TUNEL assays using transgenic expression of CifA mutants alone or together with intact CifB in the testes. Similar to individual CifA mutant expressing lines, there were very few or no sperm bundles with DNA damage in CifA mutant lines upon the individual (**Fig. S5A**) or dual expression with CifB (**Fig. S5D**; p>0.05, Kruskal-Wallis test, **Table S2**). This result indicated that CifA protein mutants inhibit the *in situ* DNA cleavage activity mediated by CifB because dual expression of intact CifAB cleaved spermatid DNA at the canoe stage (**Fig. 3**). Inhibition of such activity could occur because of mutated CifA residues hampering proper CifB binding, which may impact downstream spermatid DNA cleavage. For example, the CifA_3_ mutant residue is near the binding interface for CifA and CifB (Xiao et al., 2021). Interestingly, despite mutated residues in CifA_4_B (with substituted residues in the predicted STE domain) that did not apparently cleave spermatid DNA *in situ* (**Fig. S5D, S6**), this mutant caused strong transgenic CI (**Fig. S5F**), suggesting the CifA4 mutant sites are important for DNase activity in the canoe-stage spermatids but not CI. Similar to the CifA mutant findings, single expression of CifB mutants and dual expression of CifB mutants with wild type CifA generally ablated spermatid DNase activity (**Fig. S5B**; **S5E** p<0.0001, Kruskal-Wallis test, **Table S2**) as well as CI (**Fig. S5C, S5F, Table S3**). Overall, amino acid substitutions throughout the length of the T1 CifA and CifB proteins ablated CI and *in situ* DNase function at the canoe stage of spermiogenesis.

### QxxxY motif associates with CifA and CifB nuclease activity

T1 CifB lacks the canonical PDDEXK catalytic residues (Beckmann et al., 2017; LePage et al., 2017; Lindsey et al., 2018; Wang et al., 2022; Xiao et al.,

2021), and CifA lacks any similarity to PDDEXK nuclease domains. As such, the nuclease data above lead to a hypothesis that there may be an alternative catalytic site for nuclease activity of the *w*Mel Cif proteins. Either the PDDEXK-like domain in CifB is active, or the Cif proteins encode alternative catalytic sites or motifs to cleave DNA and RNA. Both explanations are equally viable, though several mutants within and outside the PDDEXK domains can ablate *in vitro* and *in situ* DNase function (**Fig. 1B, Fig. S1, Fig. S5**). Interestingly, T1 CifB of *w*Mel possesses a QxxxY motif within a region of predicted α-helices at the CTND (**Fig. 4A**). The QxxxY motif is characteristic of RecB-family nucleases with HsdR subunits of *Escherichia coli* that also encode a canonical PDDEXK domain (Singleton et al., 2004). Mutating Q and Y residues impairs EcoR124I DNA cleavage, possibly by stabilizing the catalytic pocket of the PDDEXK domain or modulating the binding efficacy of this domain to the DNA (Šišăkovă et al., 2008).

Notably, the QxxxY motif is also present in T4 CifB of *w*Pip characterized as a DNase, present in T3 CifB of *w*No (Adams et al., 2021; Sun et al., 2021) and missing in T1 CifB of *w*Pip previously deemed to lack nuclease function (Chen et al., 2019). Thus, it is tentatively possible that the QxxxY motif may assist CifB DNase activity since it is localized in CTND, and the sites in CTND are proposed to be involved in substrate binding or catalytic regulation (Xiao et al., 2021). Notably, CifA also possesses the QxxxY motif in one of its predicted α-helices (**Fig. 4B**), emphasizing the potential association of this motif in mediating nuclease activity. While this motif’s presence and predicted structure correlated with the *in vitro* DNase activity data of CifA and CifB homologs tested thus far, it is not possible to prescribe relative importance over the PDDEXK-like motifs since various mutants beyond these sites ablated DNase function.

## Discussion

CifA and CifB proteins are both required for CI induction in *D. melanogaster* (LePage et al., 2017; Shropshire et al., 2018; Shropshire and Bordenstein, 2019) induced by the *w*Mel strain, the same strain released in *Aedes aegypti* field trials by the World Mosquito Program (Bourtzis et al., 2014; O’Neill et al., 2018; Utarini et al., 2021). At the cellular level, both CifA and CifB target the nuclei of developing sperms in flies; and CifA harbors a nuclear localization sequence that is functionally required for nuclear entry in developing fly sperm cells and the recapitulation of CI and rescue (Kaur et al., 2022). Based on these premises, we hypothesized that the two Cif proteins metabolize nucleic acids. We thus set out to evaluate their enzymatic properties using *in vitro* and *in situ* assays. We describe five key results: (i) T1 CifB is a DNase, (ii) T1 CifA is both a DNase and RNase, (iii) *Wolbachia* and transgenic CifAB expression causes DNA damage at the canoe stage of spermiogenesis, but *in situ* DNase activity of CifB alone is insufficient to cause CI, (iv) truncations and conserved site-directed mutagenesis establish the dependency of Cif nuclease functions across the length of the proteins, and (v) a QxxxY motif in both CifA and CifB associates with observed nuclease activity. Notably, this is the first report to demonstrate nuclease functions of the CifA protein. Below, we discuss the relevance of Cif nuclease findings to the molecular bases of CI, the Host Modification model, and *cif* gene nomenclature.

Previous work showed that the T1 and T4 phylogenetic clades of CifB biochemically act as *in vitro* deubiquitinases (DUBs) and DNases, respectively. These findings led to a proposal of an enzyme-dependent gene nomenclature and mechanistic narrative of CI (Beckmann et al., 2017; Chen et al., 2019). Structural homology-based annotations reveal that DUB-encoding T1 CifBs also contain PDDEXK-like nuclease dimers (Beckmann et al., 2017; LePage and Bordenstein, 2013; Lindsey et al., 2018) - however, due to the absence of canonical catalytic residues, T1 CifBs were deemed non-nucleases (Chen et al., 2019). Notably, PDDEXK nuclease family enzymes vary substantially in sequence and structure while retaining DNase activity (Knizewski et al., 2007; Steczkiewicz et al., 2012). Indeed, CifB nuclease domains across all CI Types show homology to not only the PDDEXK family but also to the restriction endonucleases HsdR (Lindsey et al., 2018). HsdR is a subunit of type I restriction-modification enzymes (Obarska-Kosinska et al., 2008) that carries a unique and conserved QxxxY motif sequence located in the α-helix immediately C-terminal to the PDDEXK motifs (Šišăkovă et al., 2008). Mutated Q and Y residues hamper the DNA cleaving efficiency in the EcoR124I restriction enzyme, indicating their contribution to DNase activity (Šišăkovă et al., 2008). Interestingly, both CifA and CifB of *w*Mel possess the QxxxY motif in a predicted α-helix region, and in CifB, they are present in the C-terminus preceded by the PDDEXK motif. Thus, future CI studies may consider association of the QxxxY motif with DNase activity by the CifA and CifB proteins.

*In situ* assays at the canoe stage of development notably revealed spermatid DNA damage in both *Wolbachia* and dual CifAB expressing lines, thus demonstrating the Cif enzymes are responsible for metabolizing DNA. DNA damage and repair are typical features facilitating the histone-to-protamine transition during the canoe stage of spermiogenesis to tightly pack the sperm chromatin (Rathke et al., 2007). However, excessive DNA breaks result in abnormal sperm chromatin packaging that can impact male fertility and embryonic viability in mammals (Agarwal and Said, 2003; Hosen et al., 2015; Sakkas and Alvarez, 2010). *Wolbachia*-induced oxidative DNA damage was previously reported in an early spermatogenesis stage of *D. simulans* (Brennan et al., 2012) suggesting that modifications to developing sperm contribute to CI. Therefore, detection of Cif-mediated DNA cleavage of spermatids is noteworthy here as DNA damage can adversely impact Histone modifications that hampers the histone removal process from chromatin (Bannister and Kouzarides, 2011; Tjeertes et al., 2009). Indeed, the nuclear-targeting CifA and CifB proteins alter sperm chromatin integrity by increased histone retention and decreased protamine deposition during sperm development (Kaur et al., 2022). Moreover, mutant males with protamine knockouts enhance wild type *Wolbachia* CI levels (Kaur et al., 2022). Taken together, our data suggests that Cifs promote spermatid DNA damage to alter sperm genome integrity and paternally bestow a long lasting modification which underpins CI.

While dual expression of intact CifA and CifB promotes DNA damage, CifA mutants expressed with intact CifB inhibit spermatid DNA cleavage. These findings suggest nuclear-targeting CifA is central to metabolizing spermatid nucleotides, and mutations and deletions in CifA may hinder the proper binding and/or localization of CifA and CifB (Horard et al., 2022; Kaur et al., 2022; Wang et al., 2022) in testes so that they cannot access target substrates to mediate cleavage. Ablation of spermatid DNase action by dual expression of CifB and CifA_4_ (with substituted residues in the predicted Sterile transcription factor domain (STE)) is particularly notable as it induces CI. CifA’s STE domain is highly conserved across the phylogenetic Types (Lindsey et al., 2018). Crystal structure of CifA predicts this region to be disordered (Wang et al., 2022; Xiao et al., 2021). Disordered regions may facilitate post-translational modifications and regulate key protein functions (Sharma and Schiller, 2019), in this case, DNase activity that is independent of CI.

Upon individual Cif expression, we observed that CifA alone does not recapitulate *in situ* DNase activity despite its ability to function as a DNase *in vitro*.At the histone-to-protamine transition stage of spermiogenesis, *w*Mel CifA is not highly abundant, and *w*Pip CifA reportedly vanishes (Horard et al., 2022; Kaur et al., 2022). Thus, one explanation is that CifA levels are below the threshold to induce spermatid DNA nicks or it requires co-expression with the CifB binding partner (Wang et al., 2022; Xiao et al., 2021). In contrast, single CifB promotes DNA fragmentation at this stage. T4 *w*Pip CifB is also an *in vitro* DNase, which is proposed to contribute to the CI mechanism independent of CifA via DNA cleavage (Chen et al., 2019), although the host DNA substrate that T4 CifB acts on remains unknown.

In conclusion, we report previously-unrecognized nuclease functions of both Cif proteins from *w*Mel *Wolbachia* and establish a new understanding that T1 CifA is a DNase and RNase, and both T1 and T4 CifBs are DNase enzymes, which is consistent with general homology between their core nuclease domains. Based on these findings, we suggest that a durable gene nomenclature for Cifs will be grounded in evolutionary genetics and stable phylogenetic Types (T1-T5) that are less subject to shifting discoveries based on Cif enzymatic functions, as revealed here. Finally, we link *in vitro* and *in situ* enzymatic activity of the Cif proteins and highlight their impact on the nuclei of developing host sperm, a finding that is consistent with the Host Modification model of CI in this system (Kaur et al., 2022). This work highlights the significance of replicating *in vitro* enzymatic assays with *in situ* and/or *in vivo* measurements to confirm the biological functions of proteins in the arthropod host systems and their relation to the CI mechanism.

### Ideas and Speculation

Given that individual expression of *w*Mel CifA and CifB is insufficient to establish CI (LePage et al., 2017) and modify sperm genome integrity (Kaur et al., 2022), we propose that Cif-mediated spermatid DNA and/or RNA damage may be central to the incipient CI-defining sperm modifications that alter chromatin integrity (**Fig. 5**). For example, CifA RNase activity may regulate *cifB* transcripts and thus gene expression that is known to be lower than that of *cifA* (Beckmann and Fallon, 2013; LePage et al., 2017; Lindsey et al., 2018), and/or CifA RNase may degrade host mRNAs. In flies, RNA depletion in the spermatocytes can lead to sperm chromatin abnormalities and infertility (Mills et al., 2019). In addition, CifB’s DNase activity may work alone or together with CifA’s RNase or DNase functions to alter spermatid chromatin. Each of these hypotheses can impact CI (**Fig. 5**) and hypothetically, it is fair to speculate that divergent Cif homologs and host genetic backgrounds may impact the outcomes. Thus, some *Wolbachia* strains may cause CI by variations on a theme around the Cif proteins. The relative importance of CifA’s RNase activity to CI will be significant for future research.

## Materials and Methods

### 1. Fly rearing and strains

*D*. *melanogaster* stocks *y^1^w*\* (BDSC #1495), *nos*-GAL4:VP16 (BDSC #4937), and UAS transgenic (TG) lines homozygous for *cifA*, *cifB*, *cifA;B,* and *cif* mutants (LePage et al., 2017; Shropshire et al., 2020) were maintained at 12:12 light:dark at 25°C and 70% relative humidity on 50 ml of standard cornmeal- and molasses-based food medium. Lines without *Wolbachia* were previously generated (LePage et al., 2017) through tetracycline treatment for three generations. Wolb_F and Wolb_R3 primers were used to confirm symbiont presence. Virgin flies were collected and stored at room temperature.

### 2. *In vitro* Nuclease assay

Codon optimization, gene synthesis, cloning, and protein expression/purification were outsourced to GenScript Biotech (New Jersey, USA). Briefly, *cifA* and *cifB* genes were codon-optimized for translation and expression in *E. coli*, *de novo* synthesized, cloned into pGS-21a expression vector, and transformed into *E. coli* BL21 (DE3) competent cells. Cultures were grown in 1 L TB medium containing ampicillin and were incubated in 37°C at 200rpm. Once cell density reached to 1.2 O.D. at 600 nm, 0.5 mM isopropyl-b-D-thiogalactoside (IPTG) was introduced for induction at 15°C for 16 hours and then centrifuged at 8000 *g* for 20 min.

For CifA, cell pellets were lysed on ice using buffer A (50 mM Tris,150 mM NaCl, 1 mM TECP, pH 8.0) by sonication. The lysate was clarified by centrifugation at 12000 *g* for 30 min at 4°C. The clarified extracts of target CifA proteins carrying His tags at the C-terminus were filtered with a 0.45 μm filter and loaded onto Ni-nitrilotriacetic acid (NTA) agarose resin preequilibrated in buffer A. The column was then washed with buffer B (50 mM Tris, 150 mM NaCl, pH 8.0) 10 column volumes (CV) to wash away impurities from the target proteins. The protein was eluted using buffer B supplemented with 20/50/300 mM imidazole. All proteins were sterilized by passing through 0.22 μm filter before being stored in aliquots. The concentration was determined by BCA^TM^ protein assay with BSA as a standard. SDS-PAGE and Western blot were used to confirm protein purity and molecular weight (**source data files**).

For CifB, cell pellets were lysed on ice using buffer A (50 mM Tris,150 mM NaCl, pH 8.0). The lysate was clarified by centrifugation at 12000 g for 30 min at 4°C. The inclusion body pellet was solubilized in denature buffer A (7M Gu-HCl, 50 mM Tris-HCl,150 mM NaCl, pH 8.0) by sonication. The cell precipitate was spun down at 13,000 rpm for 30 min at 4°C, then the supernatant including target CifB proteins carrying His-GST tag at the N-terminus was filtered with a 0.45 μm filter and loaded onto NTA agarose resin preequilibrated in buffer A. The column was then washed with buffer B (8 M Urea, 50 mM Tris-HCl, pH 8.0) 10 CV to wash away impurities from the target proteins. The protein was eluted using buffer B supplemented with 20/50/300 mM imidazole. All proteins were sterilized by passing through 0.22 μm filter before being stored in aliquots. The concentration was determined by BCA^TM^ protein assay with BSA as a standard. SDS-PAGE and Western blot were used to confirm protein purity and molecular weight (**source data files**).

Additionally, mass spectrometry-based protein identification was performed to ensure no contaminant nuclease from the *E. coli* expression system was co-purified. Briefly, 20 μg of purified proteins (CifA, CifA_△C_, CifB_△D_, and CifB_2;△D_) were prepared for analysis using Suspension trap technology (Casiraghi et al., 2001) using the manufacturer protocol. The resulting peptides were analyzed by a 70-minute data-dependent LC-MS/MS analysis. Briefly, peptides were auto sampled onto a 200 mm by 0.1 mm (Jupiter 3 micron, 300A), self-packed analytical column coupled directly to an LTQ (ThermoFisher) using a nanoelectrospray source and resolved using an aqueous to organic gradient. A single full-scan mass spectrum followed by 5 data-dependent tandem mass spectra (MS/MS) was collected throughout the run and dynamic exclusion was enabled to minimize the acquisition of redundant spectra. The Resulting MS/MS spectra were searched via SEQUEST against a database containing the expressed proteins, an *E. coli* background proteome and reversed version for each of the entries. Identifications were filtered and collated at the protein level using Scaffold Proteome Software.

*In vitro* nuclease activity assays were performed as previously described (Chen et al., 2019). For DNase and RNase activity measures, 1μM of individual Cif proteins were incubated in a reaction buffer containing 20 mM Hepes (pH 8.0), 5 mM MgCl_2_, 2.5% sucrose, 150 mM NaCl, 0.001% Triton X-100, and 2 mM DTT with 500 nM single-stranded (ss) Cy5-labeled DNA [70-mer: Cy5-GCAATTCGATCGTTGACATCTCGCGTGCTCGGTCAATCGGCAGATGCGGAGTGAAGTTCCAACGTTCGGC-3] as previously used (Chen et al., 2019); 15nM double-stranded (ds) 154bp PCR purified fragment of *rp49* gene (Shropshire et al., 2018); or 100nM of synthetic RNA [45-mer: 5′ GGGUCAACGUGGGCAAAGAUGUCCUAGCAAGCCAGAAUUCGGCAG −3′] generated by Sigma. In reactions where CifA was co-present with CifB, 10 μM CifA was used as previously described (Chen et al., 2019). All reactions were carried out at 25°C for 120 min and quenched by adding EDTA to a final concentration of 100 mM unless otherwise noted. Samples were run in 10% TBE polyacrylamide urea gels at 180 V for 60 min. For reactions using Cy5-labeled ssDNA, gels were imaged on the Odyssey CLx imaging system. Unlabeled dsDNA and ssRNA sample gels were post-stained with GelRed (Biotium) stain and imaged with Alpha innotech imager.

### 3. *In situ* TUNEL assay

Terminal deoxynucleotidyl transferase-mediated dUTP nick-end labeling (TUNEL) assays are based on detecting single- and double-stranded DNA nicking and fragmentation, which are characteristic of apoptotic cells (Vasudevan and Ryoo, 2016). To perform the assay and detect sperm DNA fragmentation, we first set up the flies as previously described (Shropshire et al., 2020). Briefly, virginity- controlled wild type (*w*Mel+ and *w*Mel-) and TG (*nos*-Gal4:VP16) females were aged 9–11 days and mated with males (Layton et al., 2019). We used *pos*- Gal4:VP16 line since it was previously shown to drive *cifA;cifB* expression sufficient to induce near-complete embryonic death (Shropshire and Bordenstein, 2019). We collected <8 hours old wild type and TG Gal4-UAS males as the young hatched males induce strong CI levels (Shropshire et al., 2021a; Yamada et al., 2007), anesthetized on ice to stop their movement, and dissected whole testes in ice-cold 1X PBS solution. Dissected tissues were treated with 2 mM dithiothreitol to stabilize cellular proteins for 45 min at room temperature, followed by fixation in 2% paraformaldehyde on ice for 15 min. After washing in 1X PBS for 2 min, samples were permeabilized in 0.1% TritonX-100 in sodium citrate (10 mg sodium citrate, 10 μl Triton, 10 ml milliQ H_2_O) for 2 min on ice. After washing in 1X PBS for 2 min, samples were incubated with 50 μl mix of 5 μl enzyme and 45 μl labeling solution (TUNEL *In situ* Cell Death Detection Kit, TMR Red’ from Roche) for 1.5 h at 37°C in a dark humid chamber. After washing in 1X PBS for 2 min, samples were finally incubated with 50 μL of DAPI staining solution (0.2 μg/ml), mounted on a glass slide, squashed with a coverslip, and stored overnight at 4°C. Imaging was performed using green fluorescence filter excited at 488 nm laser for TUNEL and blue at 359 nm for DAPI stain at 100x magnification in All-in-one Keyence BZ-X700 fluorescence microscope. Image exposure settings were kept constant throughout the treatment groups and images were analyzed using ImageJ software. The total number of sperm bundles and TUNEL-positive bundles with damaged DNA were manually counted per testes.

### 4. Hatch rate assays

Male siblings from TUNEL assays were used to measure CI hatch rate levels as previously described (LePage et al., 2017). Briefly, males and females were paired in 8 oz bottles affixed with a grape-juice agar plate smeared with yeast. Bottles were incubated at 25°C for 24 h at which time the plates were replaced with freshly smeared plates and again stored for 24 h. Plates were then removed from bottles, and the numbers of eggs on each plate were counted. Any crosses with fewer than 25 eggs laid were discarded from the count. After another 30 h incubation at 25°C, the remaining unhatched eggs were counted. The percent of eggs hatched into larvae was calculated by dividing the number of hatched eggs by the total egg count and multiplying by 100.

### 5. *In silico* prediction analysis

Protein sequence alignment of CifB orthologs from different *Wolbachia* strains (NCBI accession number, CifB T1 *w*Mel - WP_010962721.1, T1 *w*Pip - WP_012481788.1, T4 *w*Pip - WP_007302979.1) and CifA T1 *w*Mel (WP_010962721.1) were performed using the MUSCLE plugin (Edgar, 2004) in Geneious Prime v2021.0.3 (Kearse et al., 2012). Secondary structure predictions of CifA and CifB protein sequences were made using the PSIPRED Protein Sequence Analysis Workbench program (Jones, 1999). We manually curated the presence of the QxxxY motif based on its localization within a region of predicted α-helices in the nuclease domains in CifB and throughout the length of CifA proteins.

### 6. Statistical analysis

All statistical analyses were performed using GraphPad Prism 9 software. While comparing *in situ* TUNEL data between two groups, we used a two-tailed, non-parametric Mann-Whitney *U*-test. For comparisons between more than two data sets, we used a non-parametric Kruskal-Wallis one-way analysis of variance test followed by a Dunn’s multiple correction. This allowed robust testing between all data groups while correcting for multiple test bias. For CI hatch rate assays, statistical significance was determined by Kruskal-Wallis and Dunn’s multiple correction tests. All P-values are reported in **Table S2 and S3** and raw data files related to each experiment are included in the source data files.

## Acknowledgments

We thank members of the Bordenstein lab, especially, Sarah Bordenstein, Luis Mendez, and Luis Martinez for helpful discussions during the writing process. We thank Cell & Development Biology Equipment Resources at Vanderbilt core facility for their help in using Odyssey CLx imaging system. We also thank the Proteomics Core of the Mass Spectrometry Research Center at Vanderbilt University for performing protein identification service.

## Author contributions

R. K., J.D.S., B.L., and S.R.B. designed the research; R.K. and B.L. performed research; R.K. and S.R.B. analyzed data; R.K. wrote original draft of the study. J.D.S., B.L. and S.R.B. edited and provided feedback on the paper. R.K. and S.R.B. wrote the final version of the paper.

## Supplementary figure titles and legends

**Supplementary figure 1.**
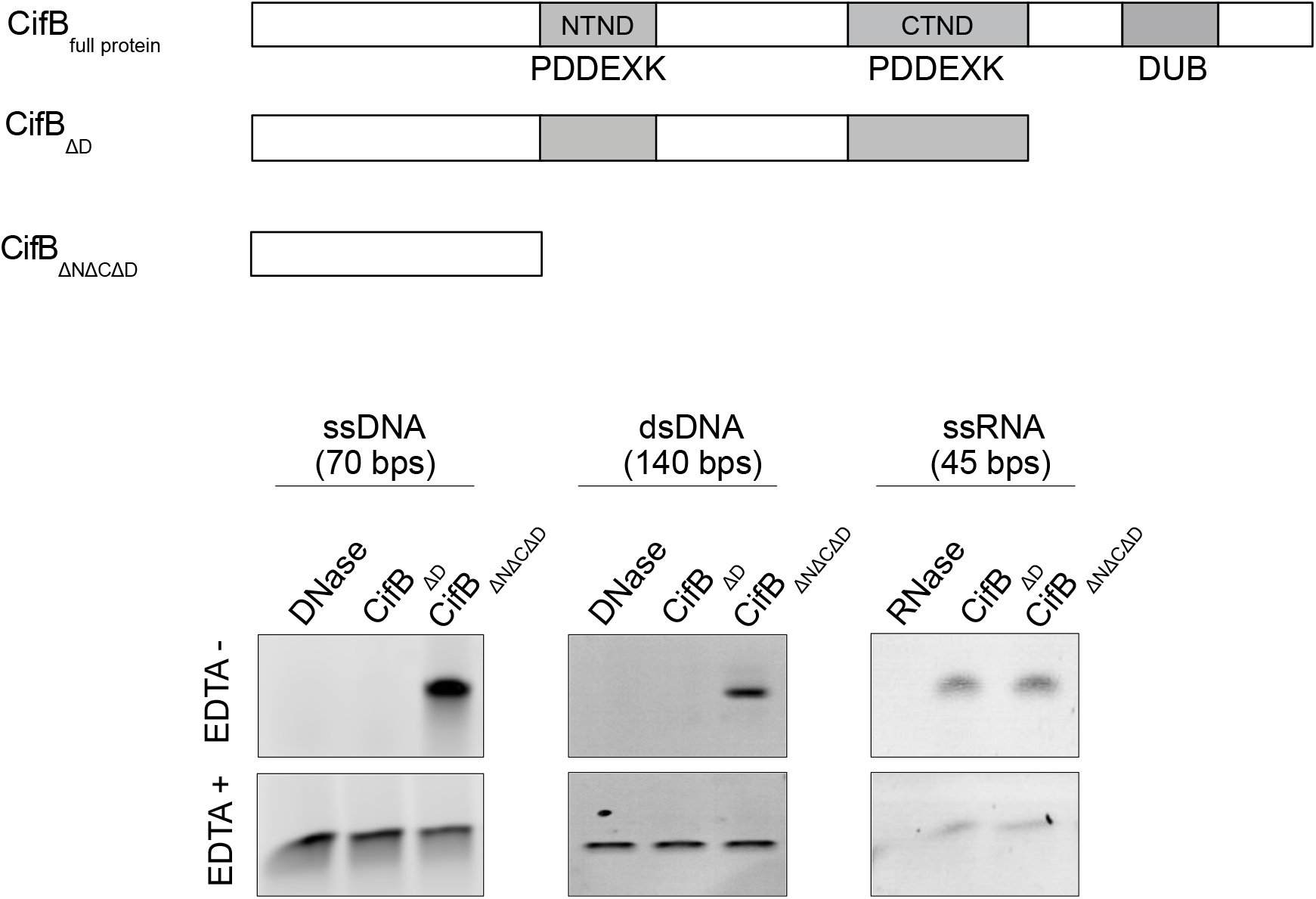
CifB amino terminus alone does not induce DNase or RNase activity. Schematic representation showing the location of truncated domains of CifB recombinant proteins used in the study. 1 μM of Cif protein was incubated with 500 nM Cy5-labeled ssDNA, 15nM dsDNA and 100nM ssRNA for 120 min. DNase and RNase enzymes were used as positive controls. To stop the reactions, EDTA was added to a 20x molar excess over Mg2+. Cy5-labeled ssDNA samples were run in a 10% polyacrylamide/TBE gel. Non-labeled dsDNA and RNA samples were run in 10% polyacrylamide/TBE gel and stained with GelRed.

**Supplementary figure 2.**
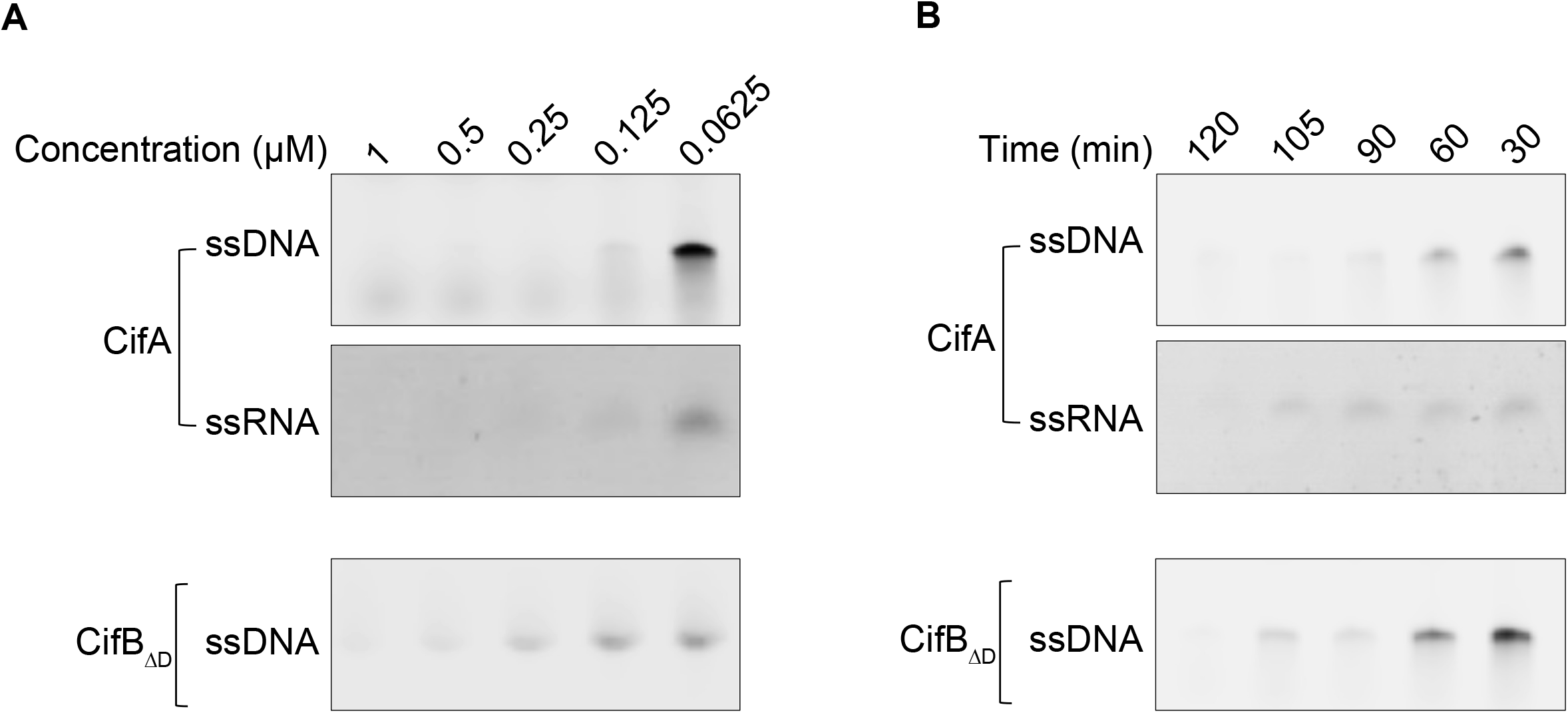
CifA and CifB are *in vitro* nucleases. (A) Dilution-based nuclease assay shows CifA’s DNase and RNase, and CifB’s DNase activities persist at higher protein concentrations and diminish with dilutions. (B) Time–course assay shows nuclease activity of Cifs diminishes at shorter incubation time points.

**Supplementary figure 3.**
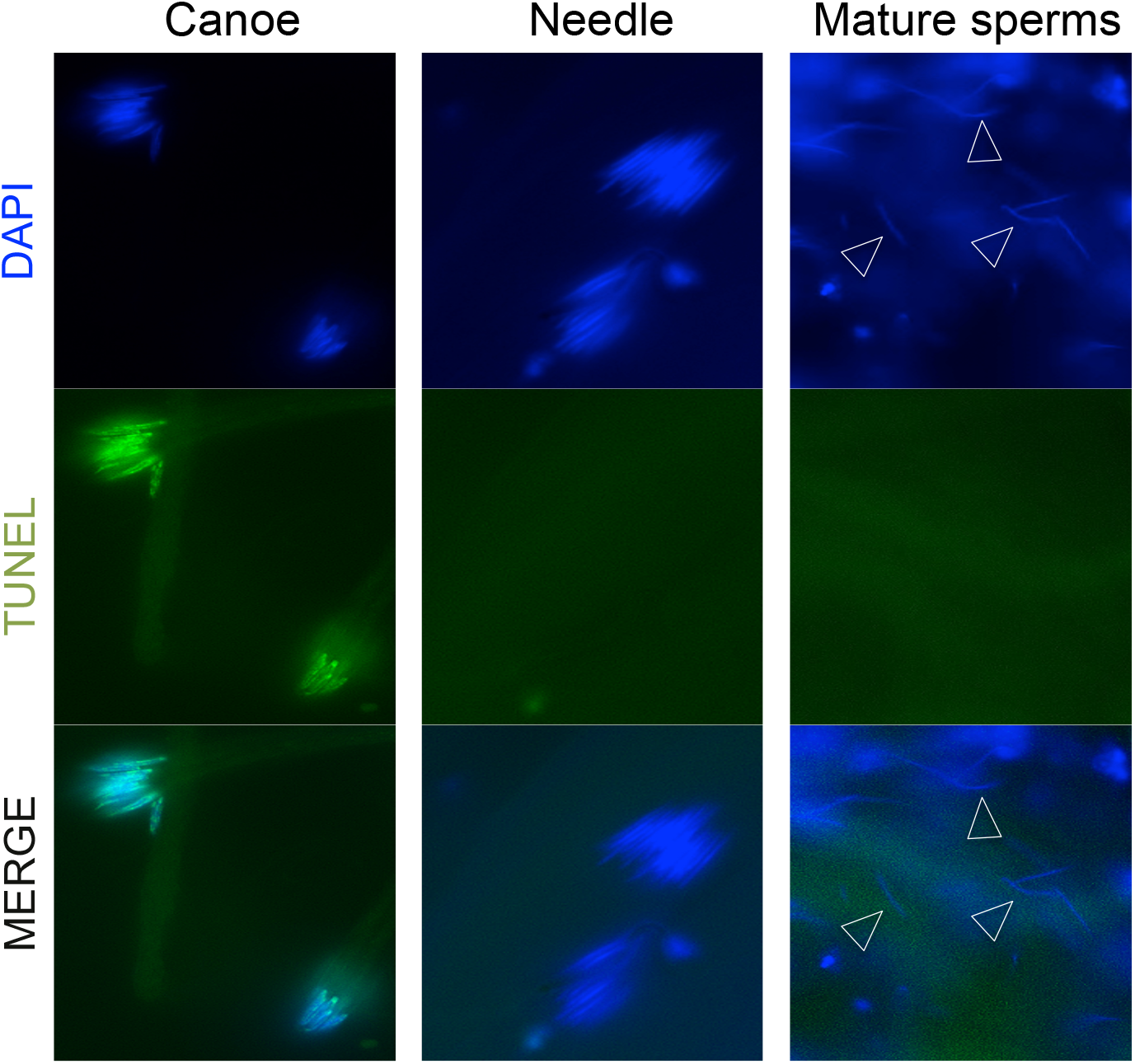
DNA breaks are detected at the canoe stage of *Drosophila* spermiogenesis. TUNEL staining on testes squashes of 0-8 hrs old males was performed to visualize DNA breaks during different stages of spermatogenesis in the same animal. DNA breaks marked by TUNEL staining are detectable only in canoe-stage spermatids, as expected (Rathke et al., 2007). During individualization, DNA breaks are no longer detectable in needle stage spermatids and the mature individualized sperms (white arrowheads).

**Supplementary figure 4.**
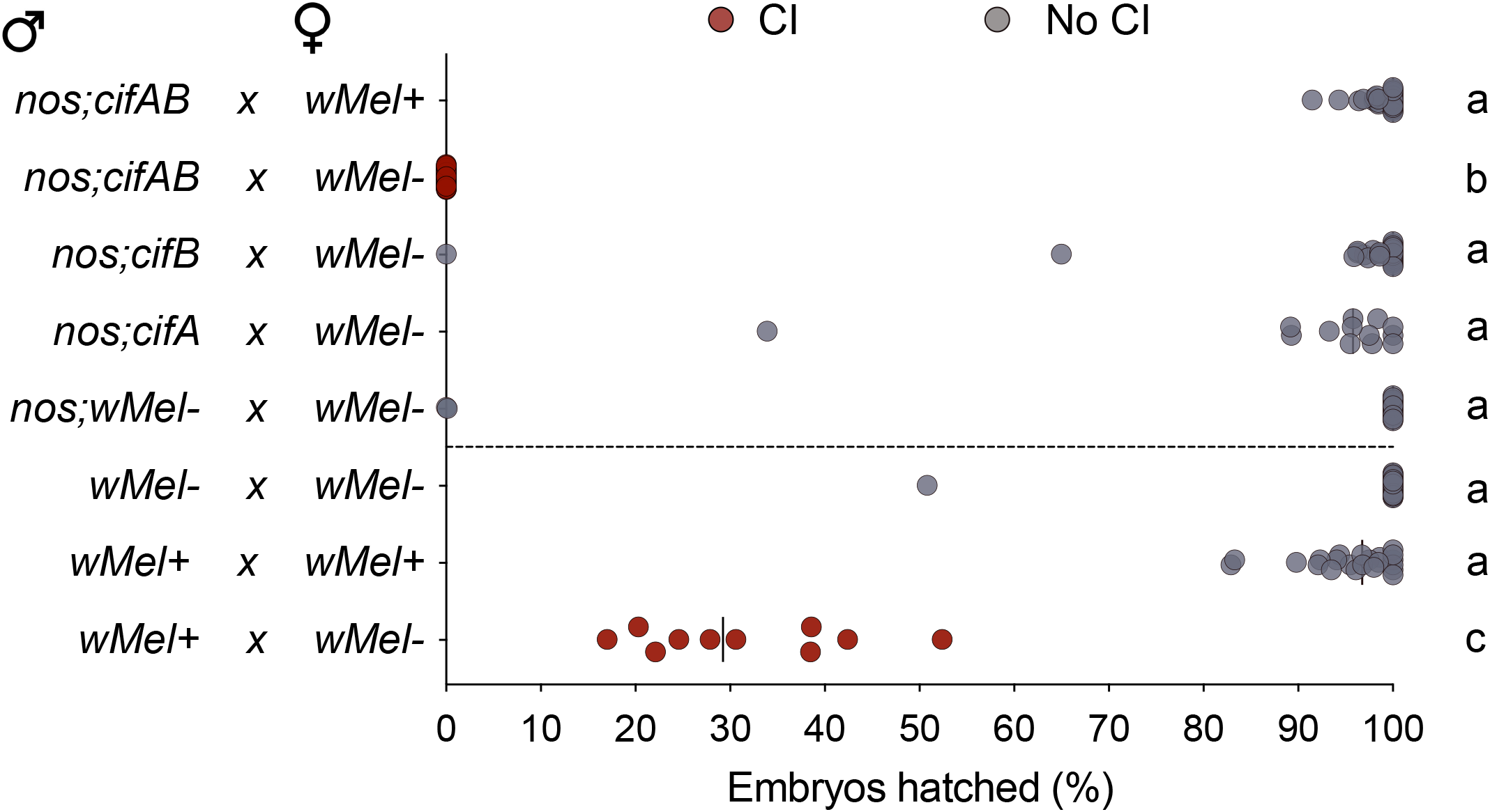
Single transgenic expression of *cifB* does not induce CI in *D. melanogaster*. Hatch rate assays were conducted to test if wild type *Wolbachia*-carrying symbiotic males and transgenic, aposymbiotic males expressing single and dual *cifA* and *cifB* genes (unfilled sex symbols) induce CI. Each dot represents the percent of embryos that hatched from a single male and female pair. Letters to the right indicate significant differences based on p = 0.05 calculated by Kruskal-Wallis and Dunn’s test for multiple comparisons between all groups. This experiment was conducted three times in parallel to the *in situ* TUNEL assay of Figure 3. P-values are reported in Table S3.

**Supplementary figure 5.**
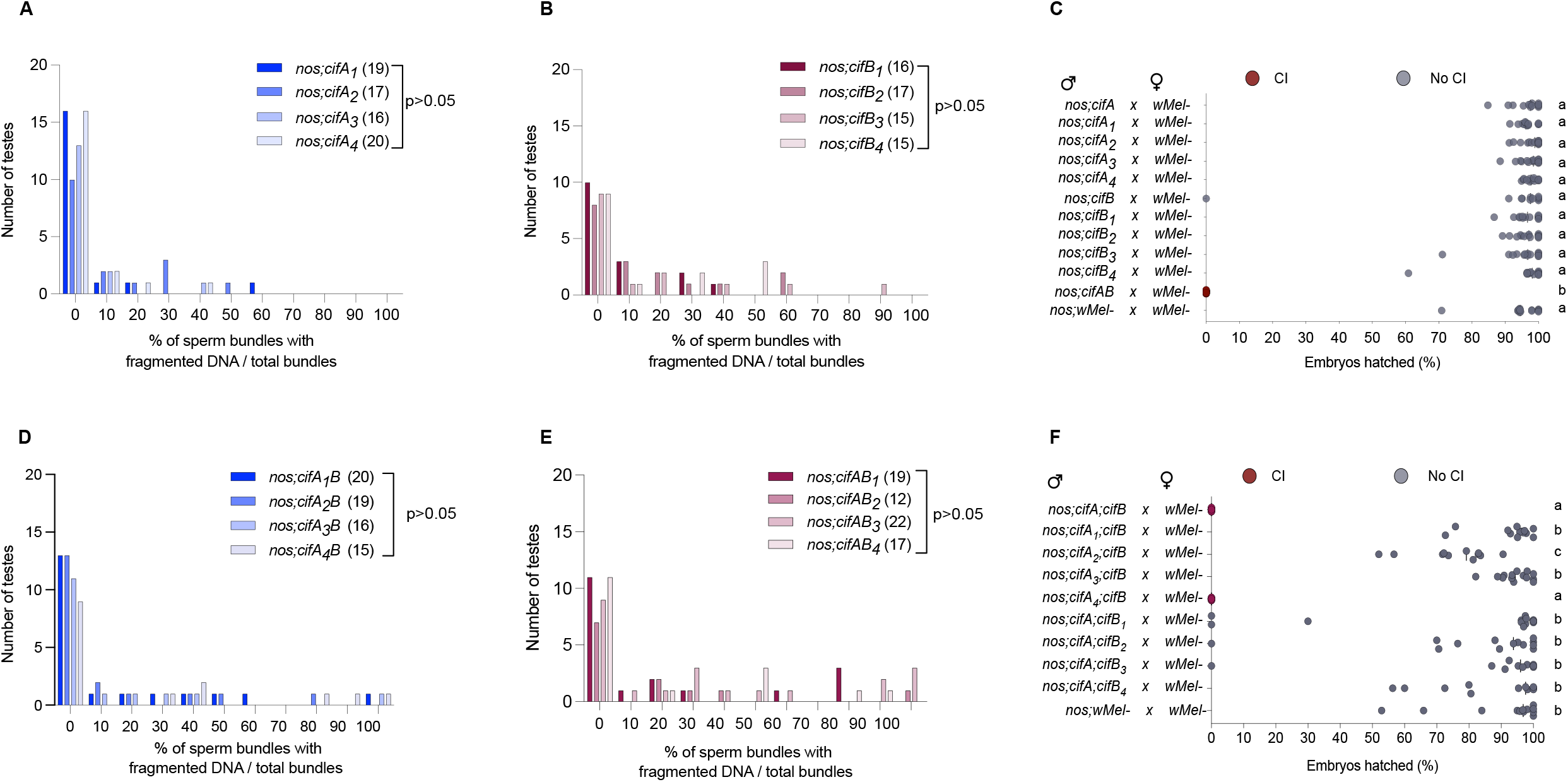
Single and dual expression of various *cifA* and *cifB* mutants ablates *in situ* nuclease activity and CI phenotype. (A, B, D, E) TUNEL assays on testes squashes of <8 hrs old males was performed to quantify sperm bundles with DNA breaks across the genotype treatment groups. Single and dual expression of *cifA* and *cifB* mutants in transgenic lines fails to induce spermatid DNA fragmentation. Total sperm bundles and those with fragmented DNA were manually counted from the images acquired. The numbers of testes investigated are shown in parentheses next to the genotype. The experiment was performed in two independent biological replicates in the same setup as experiments in Figure 3. P-value significance was calculated by Kruskal-Wallis and Dunn’s test for multiple comparisons between all groups. P-values are reported in Table S2. (C, F) Hatch rate assays were conducted to test if single and dual expression of *cifA* and *cifB* mutants can induce CI when transgenically expressed in aposymbiotic males (unfilled sex symbols). Each dot represents the percent of embryos that hatched from a single male and female pair. Letters to the right indicate significant differences based on p = 0.05 calculated by Kruskal-Wallis and Dunn’s test for multiple comparisons between all groups. The experiment was conducted twice in parallel to the TUNEL assay. P-values are reported in Table S3.

**Supplementary figure 6.**
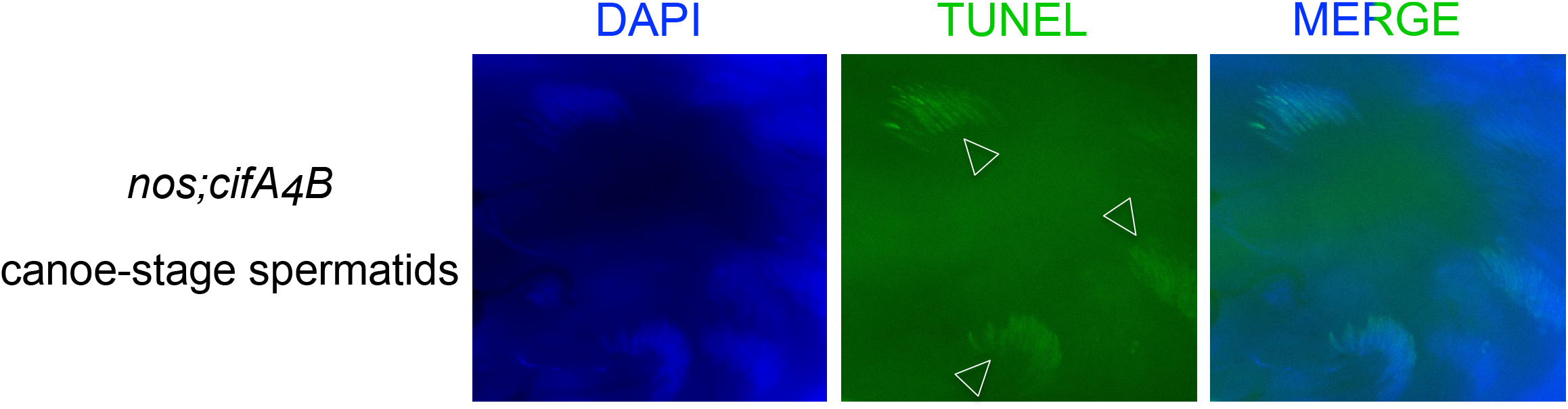
Spermatid DNase activity is ablated upon dual expression of *cifA_4_* and *cifB*. Image shows DNA nuclei (blue, DAPI) and TUNEL signal (green) in sperm bundles with less DNA breaks (empty arrow heads) compared to *cifAB* in Figure 3. The experiment was performed with two biological replicates in the same setup as that in Figure 3.

